# Frequency chasing of individual megadalton ions in an Orbitrap analyzer improves precision of analysis in single molecule mass spectrometry

**DOI:** 10.1101/2021.06.15.448530

**Authors:** Tobias P. Wörner, Konstantin Aizikov, Joost Snijder, Kyle L. Fort, Alexander A. Makarov, Albert J.R. Heck

## Abstract

To enhance the performance of charge detection mass spectrometry, we investigated the behavior of macromolecular single ions on their paths towards and within the Orbitrap analyzer. We discovered that ions in mass beyond one megadalton reach a plateau of stability and can be successfully trapped for seconds, travelling a path length of multiple kilometers, thereby enabling precise mass analysis with an effective resolution of greater than 100,000 at *m/z* 35,000. Through monitoring the frequency of individual ions, we show that these high mass ions, rather than being lost from the trap, can gradually lose residual solvent molecules and, in rare cases, a single elementary charge. Our observations highlight the importance of efficient desolvation for optimal charge detection mass spectrometry and inspired us to implement multiple improved data acquisition strategies. We demonstrate that the frequency drift of single ions due to desolvation and charge stripping can be corrected, which improves the effective ion sampling 23-fold and gives a two-fold improvement in mass precision and resolution, as demonstrated in the analysis of various viral particles.

## Introduction

Mass spectrometry (MS) offers a versatile analytical approach to characterize molecules with high precision and throughput and can be applied to molecules in mass ranging from a few to several million dalton (Da). For analysis of molecules at the higher end of this mass range, *e*.*g*., macromolecular assemblies, electrospray ionization under non-denaturing conditions (“native MS”^1^) is the method of choice to make such “elephants fly”^2^. Native MS uniquely operates under non-denaturing conditions, retaining non-covalent interactions into the gas phase. Native MS can provide unique insight into the composition, architecture and function of large biomolecular assemblies, and has been used to study, among others, ribosomal particles^3^, membrane protein complexes^4^, virus-like particles^5^ and endogenous viruses^5–7^ (also reviewed in ^8–11^).

The mass analysis of macromolecular assemblies is still less efficient than the analysis of smaller molecules (*e*.*g*., peptides, metabolites and proteins) as these large assemblies suffer from sub-optimal transmission through the mass analyzer, and often generate insufficient signal on a detector. Moreover, most mass analyzers exhibit lower resolving power at higher mass-to-charge ratio (*m/z*), which often makes it impossible to resolve sufficient features in the mass spectra of complex/heterogeneous samples. Macromolecular assemblies often represent heterogeneous samples, due to variations in composition, cargo-loading and or protein post-translational modifications. To better tackle such heterogeneous, high mass samples, single particle approaches have emerged also in mass spectrometry, circumventing the need to resolve and convolute ions signals. Two such single particle approaches are charge detection mass spectrometry (CDMS)^12,13^ and nanoelectromechanical systems-MS (NEMS-MS)^14^. In CDMS, an independent measure of the charge, *z*, is made, overcoming a major bottleneck in charge state assignments in crowded *m/z* spectra. In NEMS-MS, the ions are deposited on small resonators, and a mass for each ion is derived by detecting the resonator frequency shift caused by the deposition of the particle. Both these approaches have provided impressive results for highly heterogeneous samples like exosomes or viral particles at up to 100 MDa ^15–17^.

More recently, two groups have in parallel demonstrated CDMS on an Orbitrap™-based mass spectrometer, by utilizing the linear dependency between the ions’ charge, the induced imaging current of the time domain signal, and the resulting peak height in the Fourier transformed mass spectra^18,19^. CDMS using Orbitrap analyzer offers the possibility to investigate in depth the behavior of single macromolecular ions within the mass analyzer, which is exciting as most of our current knowledge about ion behavior in the orbital trap is derived from ensemble measurements of smaller and/or denatured particles^20^. Here we demonstrate that such knowledge is beneficial for further progress in native MS and CDMS applications on samples of ultra-high mass.

Of particular impact on the sensitivity and quality of Orbitrap mass spectra is loss of ion signals during the mass analysis period, whereby ions are lost due to unstable trajectories arising from metastable decay, space-charge effects or via collisions with background gas molecules. A single collision of a low *m/z* (below ∼2000) denatured protein or peptide ion with background gas typically results in the complete loss of the ion within the orbital trap^20–22^, making ultra-high vacuum (*i*.*e*. < 1e-9 - 1e-10 mbar) conditions a necessity. For lower *m/z* ions, the energy transfer in such collisions is high enough to cause fragmentation and expel the ions from their coherent motion, causing them to go into unstable orbits^21^. The probability of ions colliding with neutral gas molecules depends on the pressure in the trap, collisional cross section (CCS) and the travelled distance during the trapping event. The travelled distance largely depends on the trap dimensions, recorded transient time and oscillation frequencies of the analyte ions.

Compared to standard conditions typically used for applications such as proteomics and metabolomics, macromolecular analytes in native MS acquire substantial fewer charges during the electrospray ionization process. The resulting high *m/z* ions are less efficiently controlled by the ion guides due to their high inertia, making it much more challenging to achieve sufficient transmission. Efficient transmission of high *m/z* ions in native MS can be improved via additional collisional cooling in the front end of the analyzer and in the C-trap^23^. However, such approaches may also lead, through leakage, to an undesired increase of pressure in the Orbitrap analyzer. In theory, this elevated pressure could be a major bottleneck for maintaining stable ion trajectories over longer transient times, especially considering that macromolecular ions also have higher collisional cross-sections. Nonetheless, the first Orbitrap-based single particle MS studies revealed that surprisingly high stabilities were observed for high mass ions, even when recording transients of 1 second ^19^.

Here, we investigate the extraordinary stability of these macromolecular ions in the Orbitrap analyzer. We find that such ions can easily survive multi-second long transients, notwithstanding the fact that they experience a multitude of collisions with background gas molecules during that time. The energy deposited by these collisions does not cause fragmentation, as is the case for small and denatured ions but does lead to gradual solvent loss accompanied in rare instances by charge stripping events. These events are the underlying processes of temporally unstable frequencies, resulting in Fourier transform artifacts (upon processing of the transient by the enhanced Fourier Transform (eFT) technique) impairing the performance of Orbitrap-based CDMS.

To characterize the behavior of each single ion we introduce a “frequency chasing” method, allowing us to extend the transient recording times even further while minimizing FT artifacts. Such long transient recording additionally enabled the first ever experimental observation of the radial ion motion in an Orbitrap analyzer. Moreover, based on our observations of the behavior of single ions within the mass analyzer, we introduce multiple improved data acquisition strategies for Orbitrap-based CDMS. In combination, these improvements lead to a substantial improvement in effective ion sampling, resulting in better statistics and, by harnessing longer transient data, an almost two-fold increase in mass resolution.

Although still in its infancy, we foresee that ultrasensitive analysis of macromolecular assemblies by single particle Orbitrap-based CDMS will develop into an essential research tool. Research in CDMS to date has focused primarily on virus analysis. Viruses are generally of interest, but also more frequently used as gene-delivery vectors (especially Adenoviruses and AAV viruses). Therefore, the improvements in Orbitrap-based CDMS proposed here will benefit applications in virology, biotechnology and gene-delivery. As such analyses can now be performed on instruments, widely used already within the research community, it will likely open up CDMS-based single particle analysis to a broader user base.

## Results

### In FT-MS mass resolution and signal-to-noise scale with transient recording times

In image current based detection, as employed in Fourier transform mass spectrometry, prolonged recording of stably oscillating ions should lead to a higher mass resolution^24^. Whereas in conventional ensemble-based native MS the effective resolution is a convoluted result of overlapping isotopes, solvent adducts and true resolving power of the mass analyzer^24^, single ion detection enables the precise assessment of the instrument’s performance by analyzing individual ion signals, avoiding the detrimental effects of isotope interferences. For truly stable oscillating ions, there should be a linear trend of increased mass resolution with longer transient times. Here we explored whether this is indeed the case. We analyzed single ions of the 3-4 MDa hepatitis B virus (HBV) capsids and ∼9.4 MDa Flock House Virus (FHV) in the Orbitrap analyzer at several transient recording times between 128 ms and 4,096 ms under minimized pressure settings (Figure 1).

**Figure 1:**
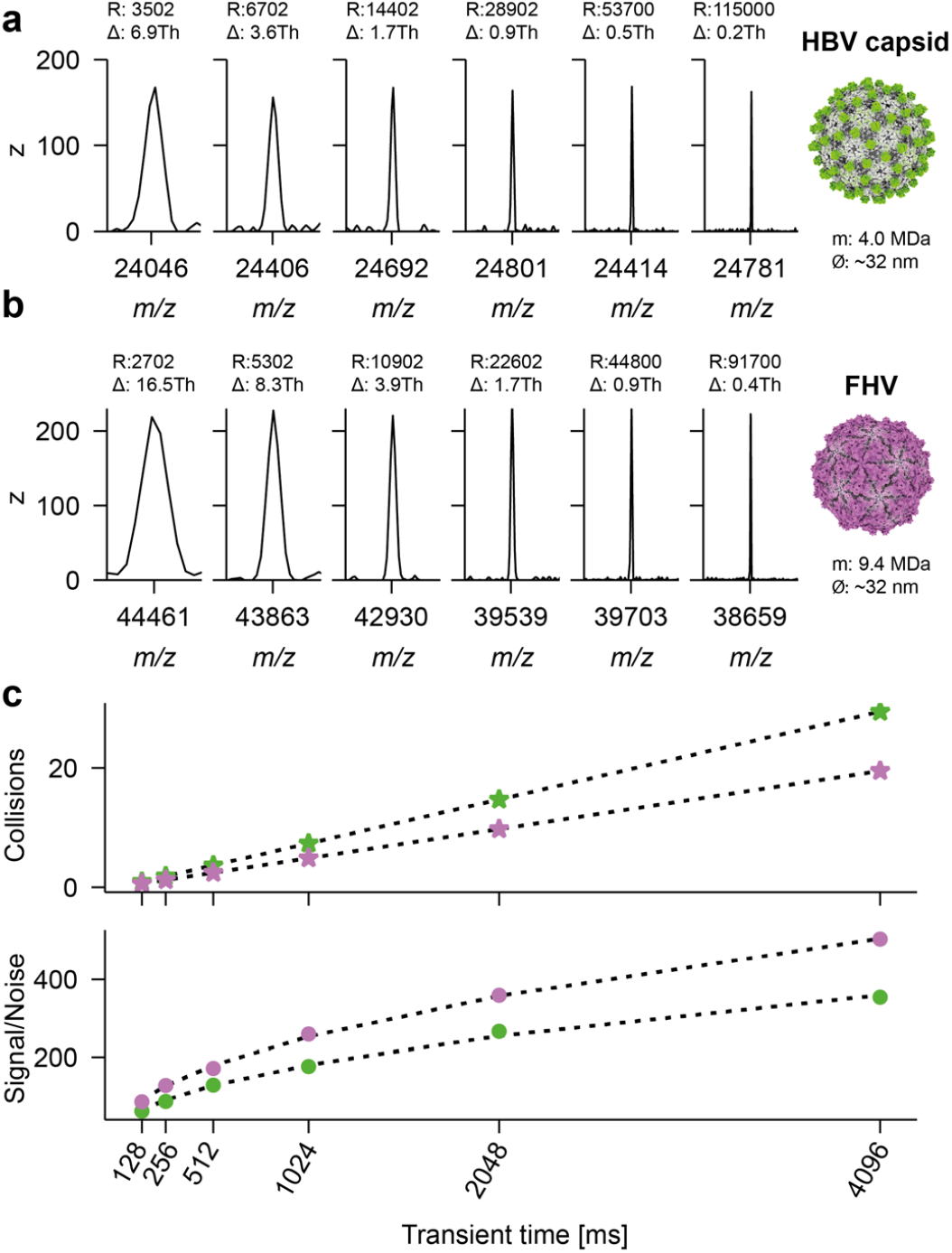
Mass resolution and signal-to-noise in Orbitrap-based CDMS scale with the transient recording time. **a**, Ion signals of individual ions for HBV at increasing transient times of 128, 256, 512, 1024, 2048 and 4096 ms (left to right), with a mass resolution for the 4 MDa particles extending from ∼3000 at 128 ms to above 100,000 at 4096 ms. **b**, as in **a** signals of individual ions of the 9.4 MDa FHV particles, with R approaching 100,000 at 4096 ms. **c**, Average number of collisions the ions experience with background gas (Xenon) during the transient time (top), with the green dots showing data for HBV and the purple dots for FHV. Although the number of collisions increases linearly, nearly all high mass ions seem to survive as evidenced by the observed resolution and signal-to-noise (bottom) that scale exactly as expected with transient time, reaching a value of ∼400 for the 9.4 MDa FHV particles at 4096 ms. Therefore, for high mass ions (Mw > 1 MDa) even longer transient times would lead to even higher resolution and signal-to-noise levels.

For all of these particles, we observed the expected linear increase in resolution (R) with transient recording time, with R increasing from ∼3,000 at 128 ms to around 100,000 at 4,096 ms. This level of mass resolution is unprecedented for particles of this size and makes it theoretically possible to resolve mass differences of less than 9 ppm. This corresponds to a Δmass of 40 Da and 90 Da for HBV and FHV, respectively, or around 0.001% of the mass of the analyte.

These multi-second long stable trajectories of individual ions are quite remarkable, as the average number of collisions they experience also scales linearly with the transient length. It can be estimated that these macromolecular particles experience around 30 collisions on average with background gas molecules during a ∼4s transient time (see Figure 1c). For individual ions, signal-to-noise (S/N) can be used as a proxy for ion survival as it will be reduced when the ion is lost or absent from later segments of the recorded transient. The plotted S/N in Figure 1c follows, for both the HBV and FHV particles, the theoretically predicted square root dependency on the transient duration, confirming that these high mass ions survive and are stable over the whole transient time, despite undergoing numerous collisions.

### A plateau of stability is reached in the Orbitrap analyzer for megadalton particles

To further understand the distinctive stability of megadalton particles compared to smaller protein ions, we analyzed single ions of a series of successive IgG1 oligomers (termed IgG1-RGY). The engineered IgG1-RGY variants have the merit of forming in solution oligomers from monomers up to hexamers, yielding ions across a wide mass range within one sample (with molecular weights of ∼150, 300, 450, 600 and 900 kDa)^25,26^.

Furthermore, we included in our analysis an even higher mass particle, the 3 MDa engineered nanocage AaLS-neg^27,28^. The binned detected centroids for both IgG1-RGY and AaLS-neg are depicted in Figure 2a. We used a segmented FT analysis^29–33^ of the recorded transients, dividing them in two halves, to estimate ion survival ratios in a semi-quantitative manner (see Figure 2b).

**Figure 2:**
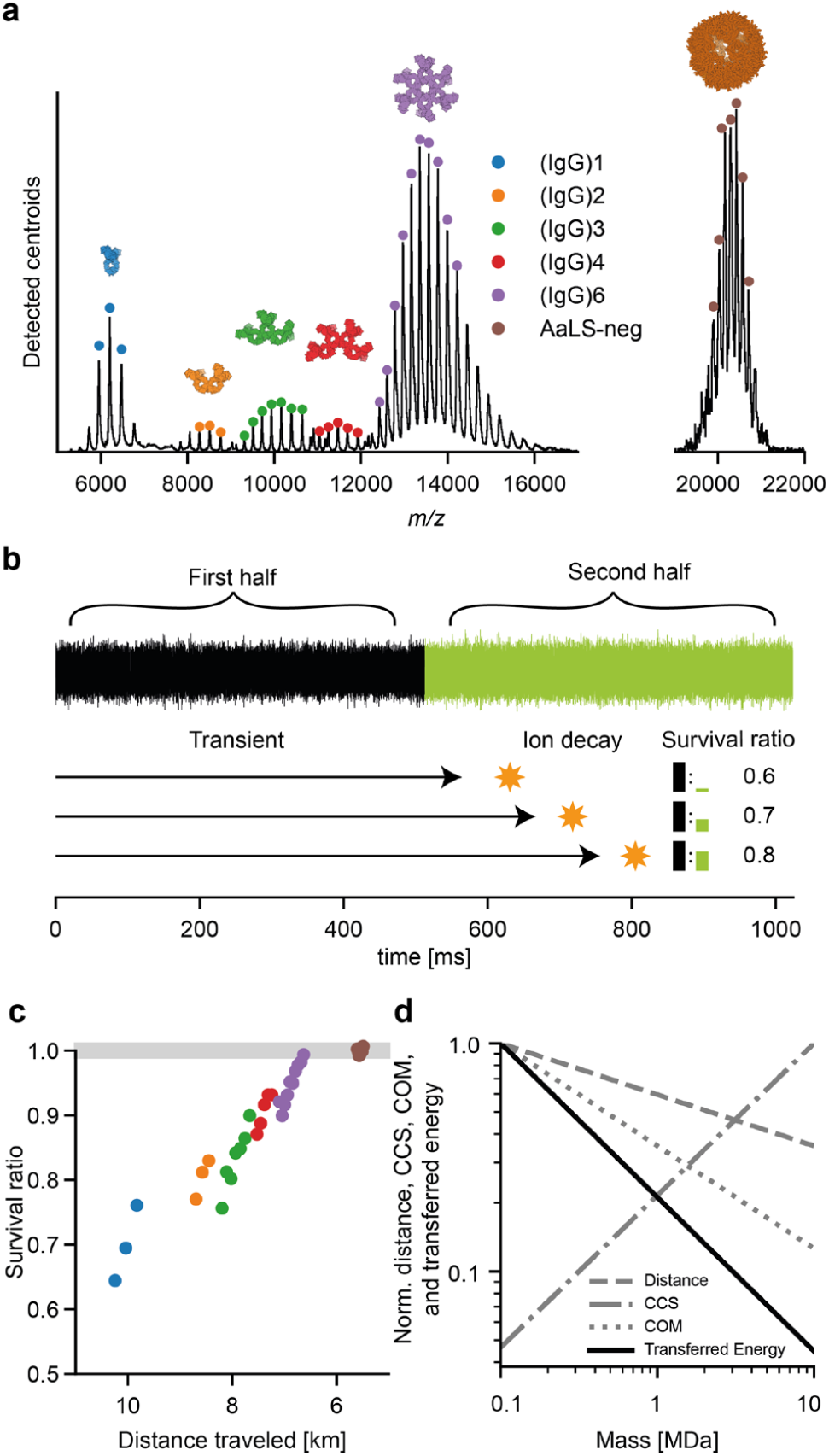
Exquisite high stability for megadalton particles in Orbitrap based native MS. **a**, Native MS spectrum constructed from binned detected centroids for the IgG1-RGY oligomers (i.e. IgG, (IgG)_*2*_, (IgG)_*3*_, (IgG)_4_, (IgG)_6_) and the nanocontainer AaLS-neg, both recorded at similar settings. Dots indicate charge states selected for survivability analysis. **b**, Comparing ion intensity between the first and second half of the transient is used as a proxy for ion survivability. This process is illustrated on ions which decay during different stages of the second half of the transient resulting in varying ratios for the detected ion signals (black and green bars). **c**, Comparison of ion survival ratio for different charge states of different IgG oligomers and AaLS-neg using a color code as in **a. d**, Favorable scaling of travelled distance, COM and CCS with increasing mass result in sharply reduced energy transfer per surface area, responsible for the plateau of stability reached for megadalton particles. Values on the vertical axis are shown relative to those for a hypothetical particle with a mass of 100 kDa

The ratios in signal between the first and second segments, here termed “survival ratio”, are presented in Figure 2c, where we converted the respective *m/z* positions to distance travelled over the ∼1s transient. This analysis evidently shows that a high percentage of the monomeric IgG1 ions (∼30%) do not survive for the entire 1s transient time, whereas for the higher IgG1 oligomers the survival ratio increases substantially and continually. We also observe improved survival for lower charge state ions within distinct oligomers and a general trend of improved survival for larger oligomers at the same travelled distance. The differences between charge states of the same oligomer are driven not only by the increased path length for higher charge states, as the collision probability scales linearly with the travelled distance, but also by the change in transferred center-of-mass (COM) energy which scales inversely with the *m/z* ratio. The differences in survival between oligomer species at similar traveled distances (*e*.*g*., at 8.5 km traveled distance) is the combined result of an increased CCS and decreased COM energy. The favorable scaling of traveled distance (∼frequency), COM and CCS as a function of mass results in a sharply decreased amount of transferred energy per surface area for high mass samples, as illustrated in Figure 2d. This, in combination with the additional degrees of freedom for energy redistribution, are most likely the main driving forces for the higher survival rate of the larger particles. Interestingly, these latter particles seem not to show any sign of decay in spite of the multiple collisions they experience over the course of the transient time.

### Gradual solvent loss results in a frequency drift of high mass ions in the Orbitrap analyzer

Above we demonstrated that single megadalton particles can be stably trapped successfully for durations as long as several seconds, allowing astonishing resolution and precision in mass analysis. Still, while analyzing signals of thousands of single ions, we also recorded single-ion signals that deviated from the typical Gaussian peak shape, and for a few of them extensive peak splitting was observed after Fourier transform employing the eFT approach^19^.

We hypothesized that this processing artifact is indicative of other gas phase mechanisms especially affecting high mass ions and could put constrains on the use of longer transients. Consequently, we decided to dissect the recorded transients in much shorter segments to monitor in time for each recorded single ion perturbations. We term this approach “frequency chasing” as depicted in Supplementary Figure 1.

Examples of such segmented FT analyses, whereby we tracked individual AaLS-neg ions over a ∼1s transient, are shown alongside the recorded eFT mass spectrum in Figure 3a. The displayed scan is contained in the larger dataset analyzed in Supplementary Figure 2 and highlights the three different categories of peaks we observe consistently in eFT mass spectra of individual ions. These are i) symmetric peaks without distinct satellite signals, ii) split peaks with satellite signals at lower *m/z*, and iii) a clear doublet of peaks with a matching peak at a later point in time shifted to a substantially higher *m/z* (see Figure 3). Fortuitously, especially at low pressures in the orbital trap, the vast majority of detected single ion events fall into category i), whereas type ii) and iii) events are exceedingly rare (also see Supplementary Figure 1 for quantification). Our frequency chasing technique enables the evaluation of the stability of single ion trajectories, and the investigation of potential decay mechanisms. Moreover, frequency chasing also allows a quantitative description of centroiding accuracy as a function of transient lengths, which we describe in more detail in the Supplementary Note and Supplementary Figure 3.

**Figure 3:**
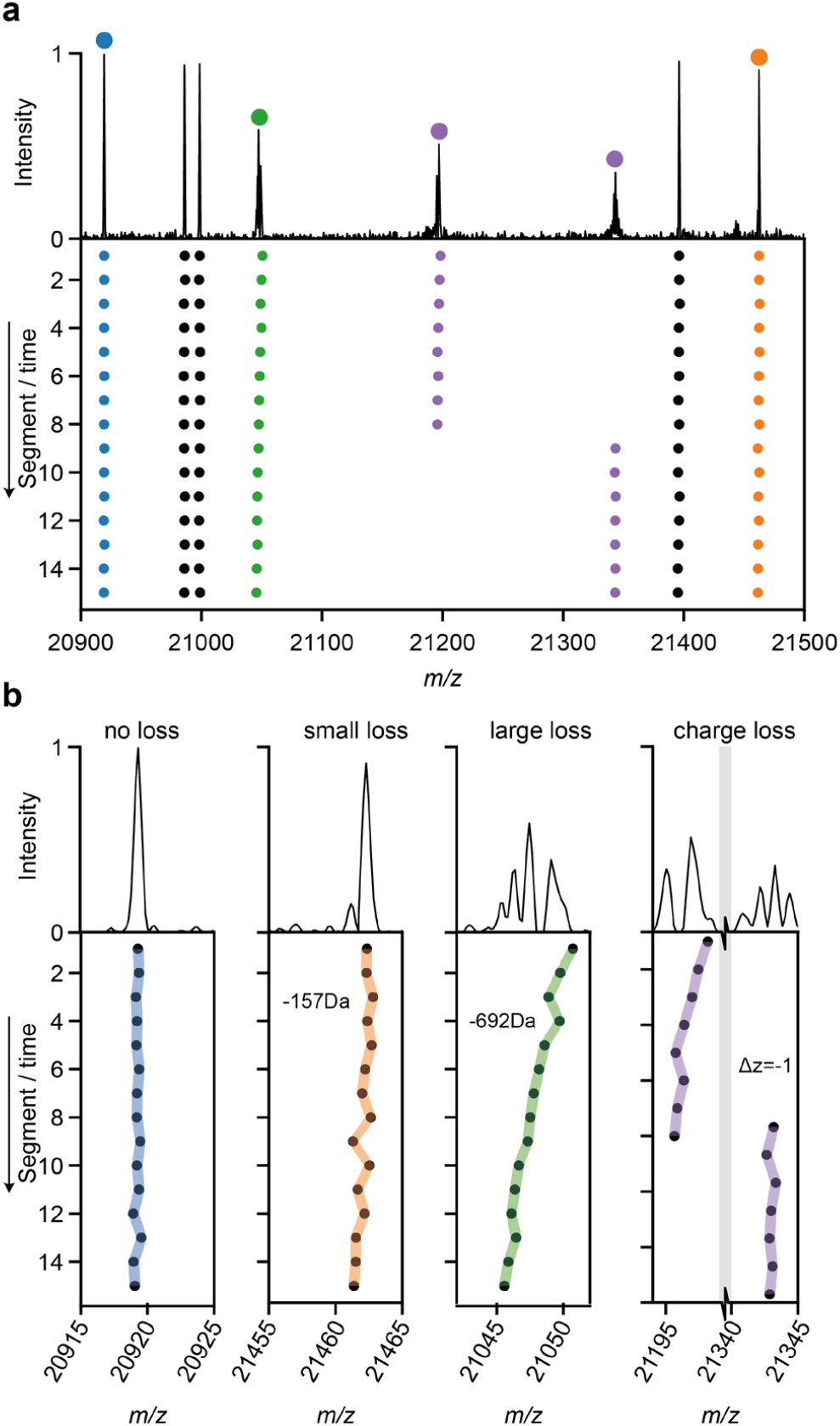
Distinctive behavior types of single ions within the Orbitrap analyzer. **a**, Single particle mass spectrum with 4 individual AaLS-neg ions highlighted (color coded). The 1s transients were divided into 15 segments and separately analyzed. The frequency and thus *m/z* of each color coded ion over these 15 segments was chased (bottom). **b**, Most single ions displayed no shifts and were stable in frequency and *m/z* (CI 95 of centroiding is shown as colored area) over the whole transient (annotated by “no loss”). A smaller percentage of the ions displayed gradual frequency shifts, and thus *m/z* shifts, due to a gradual increase in desolvation (annotated by “small to large loss”). A few ions experienced a quantized jump in frequency and thus *m/z* (gray bar indicates broken x-axis), to higher *m/z*, which is attributed to events whereby the ion loses a single charge.

The symmetric type i) peaks are completely stable in frequency within the error margin of one FT bin (denoted here as “no loss ions”). Type ii) peaks show a gradual variable shift, to lower *m/z*, hinting at a gradual consecutive loss of small solvent molecules. These mass losses result in frequency shifts, and when these exceed one FT bin, eFT processing will produce the observed split peaks with satellite signals at lower *m/z*. The type iii) peaks, which usually appear as a doublet, exhibit a sudden jump in frequency (appearing at higher *m/z*) corresponding exactly to the shift of -1 charge on an ion of near-identical mass, indicating that they are caused by a single charge loss event. Notably, these rare events allow a direct charge and mass assessment ^34^. The resulting calculated charge state, after the charge loss event represented in Figure 3 is 144.07±0.16 (±error propagated from centroiding accuracy), which is a near integer value as expected. From this value of *z* we can determine directly an accurate mass of 3,073,275±29 Da (±centroiding accuracy) for this particular single ion observed at *m/z* 21343.1±0.2.

We also monitored the frequency drift behavior of a large number of individual ions using our frequency chasing approach (see also Supplementary Figure 1) over a wider pressure range to investigate the underlying gas phase processes that govern the observed desolvation and charge loss events. Figure 4a shows the cumulative loss of individual ions for each segment with respect to the first segment for a large dataset of single ions recordings. This demonstrates a continual decrease in solvent loss over the duration of the transient, especially in the later segments. We modeled the observed trend of frequency shifts as co-occurring neutral loss processes. The first one, denoted “linear” solvent loss, occurs with a constant rate of collisions per travelled distance in the Orbitrap analyzer. These collisions are the underlying cause for the ion decay of smaller and denatured proteins as earlier described by Brodbelt *et al*.^21^. The second effect presents itself as an “exponential” solvent loss over the transient segments, in line with a suddenly activated system, which attenuates its energy gradually by solvent loss. This activation is most likely happening when the ions are injected from the C-trap into the Orbitrap mass analyzer (see Figure 4b). Although the traveled distance is short, in comparison with the several kilometers travelled during Orbitrap transient acquisition, the pressure in that region is more than a million times higher (∼10^−3^ mbar) relative to orbital trapping conditions, and ions are accelerated by approximately 1kV. We successfully simulated the solvent loss for all pressure settings using a combination of a linear and exponential decay function (with r^2^ between 0.998-0.9998). The value used for the “linear” and “exponential” solvent loss terms scale, as anticipated, linearly with the ultra-high vacuum pressure readouts in the mass spectrometer, which in turn scales linearly with the HV pressure gauge readout close to the C-trap (see Supplementary Figure 2).

**Figure 4:**
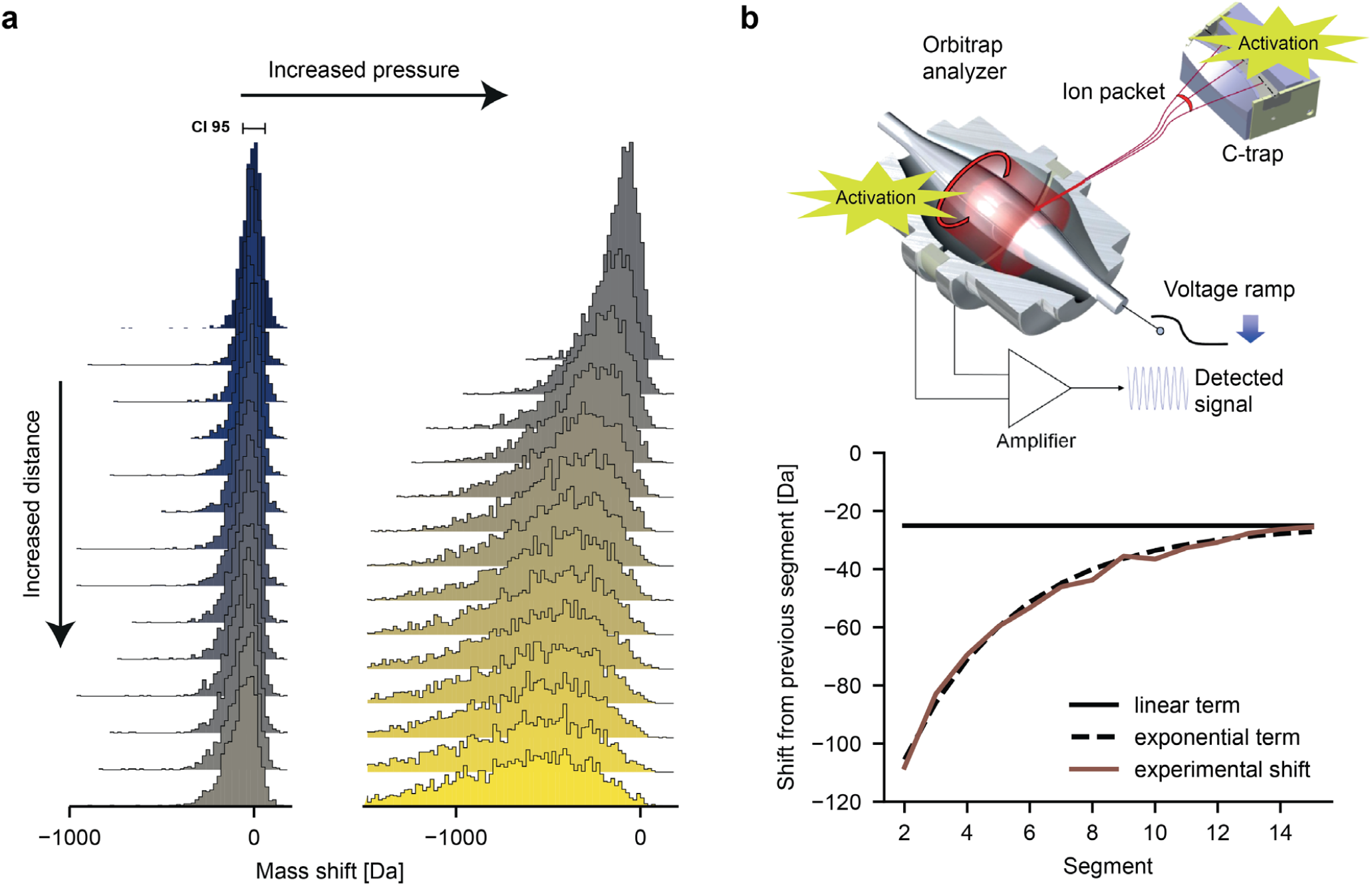
High mass ion activation and decay within the Orbitrap mass analyzer. **a**, Distribution of the observed total neutral loss for individual ions during their transient (top to bottom) at lower and higher pressure readouts (2.6*10^−10^ and 8*10^−10^ mbar, respectively). **b**, Schematic of an Orbitrap mass analyzer highlighting regions where ion activation can occur (top). The plot (bottom) depicts the average number of neutral losses relative to the previous segment under the high-pressure conditions. The black solid line indicates the constant loss rate as expected from an activation during Orbitrap detection. By adding a decaying term for the neutral loss, in line with a sudden activation in the C-trap during injection, the experimental loss can be described quite precisely. This analysis clearly reveals the dual nature of the activation process for megadalton particles.

### Optimizing Orbitrap-based charge detection mass spectrometry of megadalton particles

The core principle of Orbitrap-based CDMS is that single ion intensities scale linearly with the ion’s charge. Consequently, peak splitting caused by pressure dependent neutral losses represents a major bottleneck for the analysis of megadalton particles, especially considering that such analytes require a certain amount of gas present to facilitate effective transmission and desolvation. However, with the new insights into high mass ion behavior, we explored several experimental and *ad hoc* processing approaches to address temporal frequency instability as presented schematically in Figure 5a. In this figure for ions of the HBV *T*=4 capsid, the CDMS data on the top left was recorded at relatively higher pressure, a condition under which the eFT peak splitting occurs frequently, which significantly limits the sampling rate of ions when appropriate filtering of the split peaks is applied (∼1.5% signal utilization). This effect can be minimized by choosing suitable experimental conditions preventing either neutral loss or peak splitting. Since collisions with neutral gas molecules seem to be the main driver for neutral losses, optimizing the pressure settings can improve the relative amount of ion sampling by a factor of ∼23. Also, better desolvation of particles by increased HCD activation reduces neutral loss and peak splitting and thus increased ions sampling by a factor of ∼7. This is most likely caused by the larger amount of energy these remaining solvent molecules need to evaporate. Additionally, a reduced transient duration can reduce peak splitting by the combined effect that less time is spent in orbit, where the collisions and neutral losses take place, but also by the larger FT bin widths, which accommodate more frequency drift before splitting. This improves ion sampling by a factor of ∼13.

**Figure 5:**
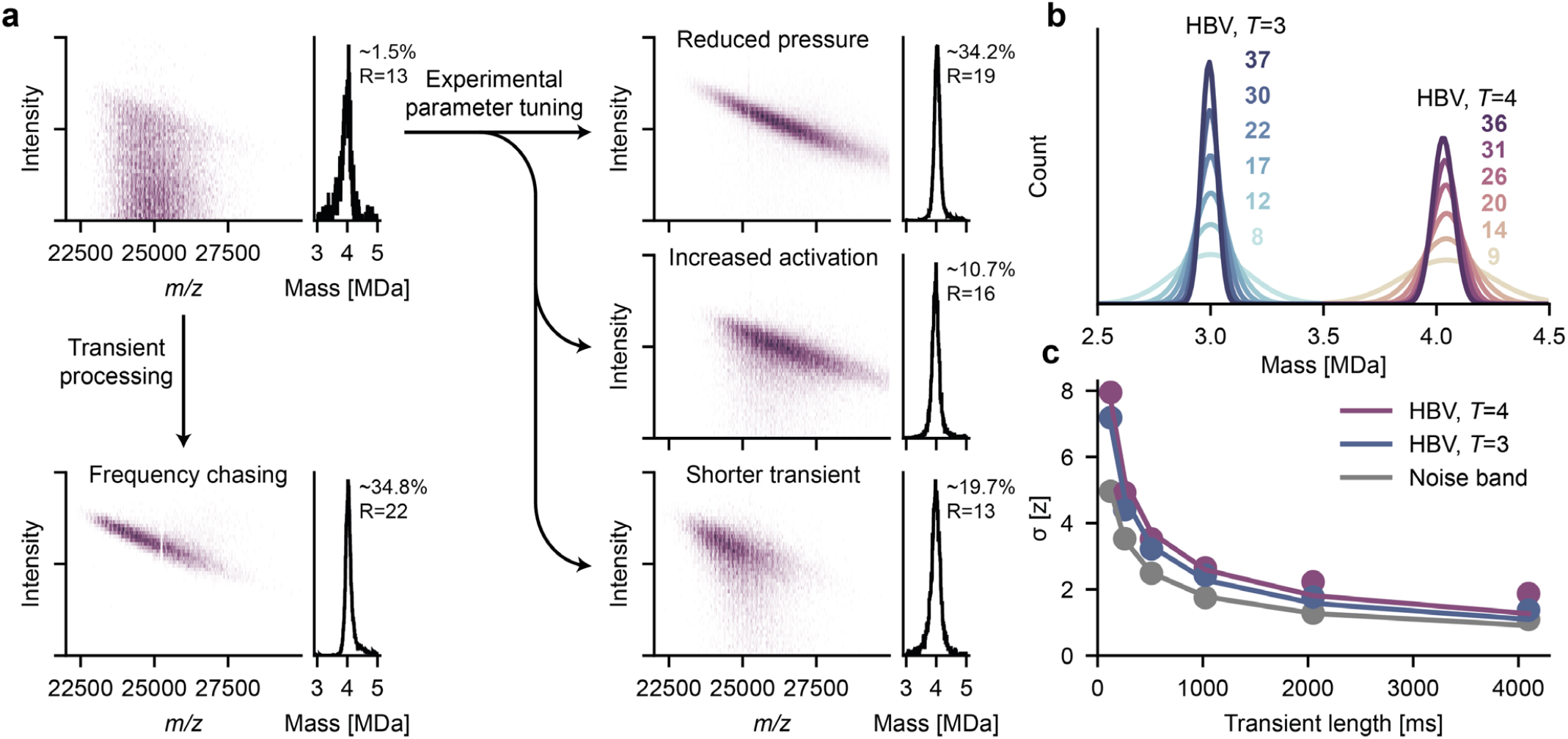
Optimizing sensitivity and resolution in Orbitrap-based single particle charge detection mass spectrometry. **a**, Overview of experimental and data processing approaches that can be exploited to reduce the number of split ion signals improving CDMS performance shown for the 4 MDa heavy HBV *T*=4 capsid. As the arrows indicate, the number of split peak occurrences can be reduced by decreasing the gas pressure (reducing the collision probability), improving the initial desolvation (lower solvent loss probability) and/or diminishing the transient time. Split peaks can additionally be diminished through frequency chasing applying drift corrections. This may overall result in a ∼23-fold increase in effective ion sampling. Each of these approaches results in a much better utilization of the acquired data and yield mass histograms with comparable resolutions. **b**, Improving mass resolution while combining longer transient times and frequency drift correction for megadalton HBV particles, *T*=3 and *T*=4 (resolution indicated in matching color). **c**, Charge resolution in single particle CDMS as a function of the transient time for HBV, *T*=3 and *T*=4 as well as of the noise band of the corresponding *m/z* region. An exponential function (σ(*t*) = *A*t*^*B*^) was fitted to the data points and follows a square root (with B: -0.52, -53, -0.49; and r^2^: 0.98, 0.99, 1.00; for HBV, *T*=4, *T*=3, and the noise band), indicating that electronic noise of preamplifier transistors is the main contributor for the observed impaired charge resolving power.

Lastly, if experimental conditions do not allow further optimization of the above-mentioned parameters, frequency drifts can be corrected using the output of our “frequency chasing” method. By taking the individual segments, like shown in Figure 2, the drift in *m/z* position is so small relative to the FT bin width that the top peak intensity stays unaffected. By subsequent averaging the intensity values of each of the segments, a similar resolution as for stable frequencies over the whole transient time can be achieved. This approach allows recovery of a large portion of the split peaks shown in Figure 5, resulting in a ∼23-fold increased utilization of the recorded ion signals.

With the use of our above-described frequency chasing methods, it is possible to accommodate the desire to record even longer transients while mitigating the effects of frequency drifts to possibly improve performance. This would not be possible with eFT-based filtering since the underlying source of such drifts (*i*.*e*. collisions with background gas) increases, while the tolerance for peak splitting (width of the FT bin) decreases with the transient time. In Figure 5b, we show the resulting mass histograms of HBV. HBV capsids can co-exist in two formations, with *T*=4 and *T*=3 symmetry, containing 240 and 180 capsid proteins, and forming particles of about 4 and 3 MDa, respectively ^35^. For the data shown in Figure 5b, we utilized increasing portions of a 4,096 ms transient while applying frequency chasing. Since the mass resolution is mainly limited by the resolution in the charge dimension, a similar trend can be seen for the spread in the intensity domain in Figure 5c. The similar scaling of the intensity spread as well as the noise band identifies electronic noise of preamplifier transistors, amplified by the relatively high capacitance of the Orbitrap electrodes, as the main contributor of non-perfect charge assignments. This noise, as well as the spread in the intensity dimension, improve following the expected square root scaling with the transient lengths closely until 2,048 ms. The deviation of the noise band is rather small and can be attributed to electronic ringing or other background processes and drift of voltages for such long transients. The larger deviations (from the theoretical square root behavior) for the actual single ion intensities compared to the noise band suggest additional artifacts, rather associated with ion behavior than pure instrument performance. Beside sample heterogeneity in the HBV capsid assemblies^36^, these artifacts are most likely caused by the wide range of kinetic energies accepted by the Orbitrap as well as eccentric, non-circular orbits, both impacting the radial distance and thus the induced image current^37^.

As an additional benefit, the recorded longer transients of highly charged particles with high S/N enables us to observe and detect the radial frequencies through their modulation of the axial ion frequency, experimentally validating for the first time their earlier theoretical description^38^.

To illustrate this, in Fig 6 we depict two instances of CDMS data on single ions where we highlight the corresponding regions at higher and lower *m/z*, displaying a stable as well as an unstable radial frequency. Notably, the resulting shape of the frequency modulations mirror themselves in the low and high *m/z* region verifying that the two modulations are caused by the same radial oscillation (see Supplementary Figure 4 for a similar observation of radial frequency modulations for single ions of GroEL and FHV and the Supplementary Note for further theoretical and experimental description of these radial oscillations).

**Figure 6:**
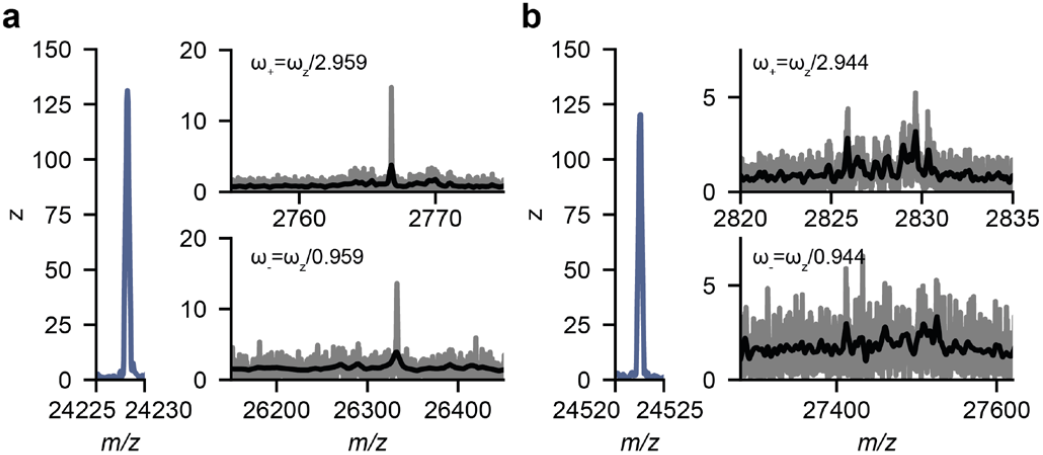
Detecting radial ion motion in an Orbitrap analyzer. **a**,**b**, show the detected peaks for two individual instances of HBV *T*=3 single ions. On the left the peak at the standard axial frequency is displayed (shown in the *m/z* domain). On the signals originating from the radial frequency modulations, ω_+_ and ω_-_are displayed, in the lower and higher *m/z* region (top and bottom). The raw data is shown in gray, the smoothed signal in black. The ion in **a** illustrates a highly stable radial frequency, whereas the single ion in **b** represents a non-stable radial frequency e.g. caused by non-circular orbits. Note, that in **b** the signals originating from the modulation by ω_+_ and ω_-_are as expected mirror images from each other in the frequency domain.

## Discussion

The foregoing detailed analysis of the behavior of individual ions of megadalton particles within the Orbitrap analyzer, including the analysis of different length FT segments, provides valuable insights allowing us to stipulate a perspective on the future of Orbitrap-based CDMS. As CDMS has unique applications in various important research areas, including structural biology, nanomaterials, fibers, vesicles, vaccines and human gene therapy (e.g. Adenoviruses and AAV gene delivery vectors), we address the need for future optimizations towards higher resolution and experimental feasibility for which we here laid out a fundamental framework.

The observed plateau of stability for megadalton particles was unexpected and seemingly in contrast to the ion behavior observed for small and denatured ions, where one collision causes ion loss through fragmentation and ion decay. Although such collisions occur frequently when analyzing megadalton particles, the deposited energy is attenuated by gradual solvent loss rather than fragmentation, allowing ions to be retained in their orbits for several seconds and possibly even for minutes if the electronics and hardware would allow us to perform such measurements. The detailed understanding of these activation processes improved CDMS experimental designs, boosting ion sampling by a factor of more than 20 (see Figure 5). Furthermore, with the frequency chasing method presented here, it is possible to extend the transient time theoretically indefinitely while minimizing the effect of frequency drifts. This is imperative as we expect the Orbitrap mass analyzer, extrapolating from the noise band, to have the capability to resolve individual charges at 16 s (*σ*=0.5) and produce an almost perfect charge assignment at 32 s (*σ*=0.3) ^39,40^. At such high charge resolutions, it is very likely that the effect of non-circular orbits and variable ion radii on the induced image current will become apparent. With the radial frequency modulations described here, we not only shed light on to the ion motion in the otherwise hidden x/z-plane of the Orbitrap mass analyzer, but also the possibility to correct for its effect on single ion intensities.

Whereas we are currently technically limited to a maximum 4 s transient time on the Q Exactive UHMR platform, developments to overcome this limit will offer the exciting prospect of improved resolution and S/N, likely beyond the crucial tipping point of determining charge state of individual ions within tolerance of a single elementary charge at a commercially available mass spectrometer.

## Materials and Methods

### Sample preparation for native MS

The purified megadalton particles analyzed were obtained from various collaborators. FHV was provided by Elizabeth Jaworski from the Routh lab (UTMB), the AaLS-neg nanocontainer sample was provided by the group of Don Hilvert (ETH Zurich), and IgG1-RGY samples were provided by the team of Janine Schuurman at Genmab (Utrecht). HBV dimer were provided by N.R. Watts (NIH). FHV, AaLS-neg and IgG1-RGY were buffer exchanged to aqueous ammonium acetate (150 mM, pH 7.5) with several concentration/dilution rounds using Vivaspin Centrifugal concentrators (9,000*g*, 4 °C). HBV capsids were formed by diluting HBV dimers cp140 directly into aqueous ammonium acetate (150 mM, pH 7.5, capsids form within minutes). An aliquot of 1–2 μl was loaded into gold-coated borosilicate capillaries 467 (prepared in-house) for nano-ESI. Samples were analyzed on a standard Q Exactive UHMR instrument (Thermo Fisher Scientific, Bremen, Germany) ^7,41^

### Instrument setting for single particle native MS

The instrument setting required for the analysis of large assemblies such as ribosomes and FHV have already been described in detail ^7,41^. In short, ion transfer target *m/z* and detector optimization were set to ‘high *m/z*’, where radio frequency amplitudes for the injection flatapole, bent flatapole, transfer multipole and higher-energy collisional dissociation (HCD) cell were set to 700, 600 and 600 V, respectively, and detector optimization was set to ‘low *m/z*’. In-source trapping was enabled with desolvation voltages ranging between −50 and −200 V. The ion transfer optics (injection flatapole, inter-flatapole lens, bent flatapole and transfer multipole) were set to 10, 10, 4 and 4 V, respectively and values of 7, 7, 7 and 7 V were used for the analysis of the IgG-RGY samples. We used either Xenon or Nitrogen as neutral gas in the collision cell and complexes were desolvated via activation in the HCD cell. Ion transmission was attenuated by diluting and shortening the injection time until individual ions were observed. Data was acquired in a range of in between 15 minutes to 2 hours. Transients were saved by setting the corresponding tab to “save with RAW file” in the “service” section of the Tune interface.

### Transient processing

The transients were four times zero-padded (as is customary^42^ in FTMS signal processing) prior to calculating its FFT. The absolute values of the FFT (i.e. magnitude mode) spectra were converted to the *m/z* domain using the instrument calibration parameters. The reported centroids of the observed peaks were calculated via a three point least squares parabola fit.

To monitor the temporal behavior of the observed masses ^29–32^ (or more accurately, the oscillation frequencies of the ionic species in the Orbitrap analyzer) during the transient acquisition, the time domain signals were segmented into a set of overlapping windows^33^. A mass spectrum for each sub-transient was computed as described above, and the masses of the ions of interest were “chased” in time. For further details see also the Supplementary Information section.

### Ion tracing

To investigate time-resolved ion behavior it is necessary to assign centroids appearing in the different segments to the ion they are originating from. Since the resolving power is much lower for individual segments converted via magnitude FT as compared to the eFT of the whole transient, only ions which were not adjacent to other centroids within a certain distance were traced. This distance is defined by the achievable resolving power at the given segment length and *m/z* position. This step circumvents also the possibility of mis-assigning centroids from two crossing ion species. After removing all closely spaced peaks, each centroid in the first segments was matched with the closest centroid (in the *m/z* dimension) of the following segments. The matched centroids are then filtered with regard to their standard deviation in the intensity and *m/z* dimension and subjected to further analysis.

### Frequency chasing

Ion frequency drift analysis, or frequency chasing, was performed on traced and filtered segmented centroid data. In order to convert frequency drifts into neutral losses all ions were binned in *m/z* and a conventional charge state assignment was performed. Next, each centroid was assigned to a charge based on its *m/z* position and frequency drifts could be converted to neutral losses in Dalton. Cumulative neutral losses and losses per segment were calculated with respect to the calculated ion mass in the first or the previous segments, respectively.

### Ion path length, CCS and neutral collision event calculations

Ion path lengths for individual ions were calculated as the product of transient length, frequency, and average traveled distance per oscillations along the z-axis. The transient length is determined by the set instrument resolution and the frequency was approximated for a given ion following the equations: f = 0.26055 * (*m/z*\*10^−3^)^-0.5^ (MHz). The average traveled distance per turn along z (93.7425 mm) was averaged over ions with different radial distances caused by the wide range of accepted ion energies for ions of the same *m/z*.

Collisional cross sections were calculated using the formula CCS= π*(r_i_+r_n_)^2≈πr_i_^2^ with r_i_ and r_n_ being the radii of the corresponding ion and neutral gas, respectively. r_i_ were extracted by manually measuring the diameter of the corresponding PDB structure. For r_n_ we used the corresponding Van der Waals radii. The average number of gas molecules in the Orbitrap analyzer was calculated from the UHV readout of the cold cathode gauge assuming the ideal gas law. We used temperature 25° C as measured by the temperature probe on the Orbitrap block. It should be noted here that the real gas pressure inside the trap is likely about 2-fold higher than measured by the cold cathode gauge according to reference ^21^. Additionally, since the installed cold cathode gauges are calibrated based on Nitrogen we applied the appropriate correction factor if used with Xenon. The mean free path of a given molecule was calculated by the product of its CCS and the concentration of gas molecules. The average number of collision events for a given ion is the quotient of traveled distance and mean free path.

### Frequency drift corrections

Frequency drifts of ions, resulting in split eFT peaks, were recovered for charge detection mass spectrometry by utilizing segmented FT analysis of the transient. Centroids from individual segments were grouped to ions as described above. These ions were then filtered for instances with a standard deviation above 4 and 0.2 for the *m/z* and intensity dimension. For ions passing these criteria, intensities and *m/z* were averaged over the individual segments. We found that magnitude FT single ions intensities can be converted under these acquisition conditions into charges by multiplying with a factor of 175.

## Data Availability

The data that support the findings of this study are available from the corresponding author upon request.

## Acknowledgements

This research received funding through The Netherlands Organization for Scientific Research (NWO) TTW project 15575 (Structural analysis and position-resolved imaging of macromolecular structures using novel mass spectrometry–based approaches), SPI.2017.028; Spinoza Award to AJRH, and NWO Gravitation 2013 BOO, Institute for Chemical Immunology (ICI; 024.002.009) to JS. We acknowledge our long-term collaborators for providing the samples analyzed, namely the Routh lab (UTMB, USA) for FHV, the group of Hilvert (ETH Zurich, Switzerand) for the AaLS-neg nanocontainer sample, and the IgG1-RGY samples from the team of Genmab (Utrecht, NL).

## Author Contributions

T.P.W., J.S. and A.J.R.H. conceived the project, designed the experiments and wrote the paper. K.A. advised and supported on segmented transient processing. K.L.F. advised and supported on long transient acquisitions. A.A.M. advised on operation of data acquisition system and mechanisms in the Orbitrap analyzer for individual high mass ions. T.P.W. performed all experiments and processed the data. J.S. and A.J.R.H. supervised the project. T.P.W., J.S. and A.J.R.H. analyzed the results. All authors discussed the results and edited the paper.

## Supplementary Information to

### Supplementary Notes

#### Accuracy of centroid determination

Detailed investigation of the behavior of megadalton single particle ions create a unique opportunity to look into their spectral idiosyncrasies from the signal processing perspective. In particular, the data exposes limits of peak centroiding accuracy in FTMS for shorter transient as well as under the elevated noise conditions^1,2^. In the current investigation all the centroids were determined using a parabola fit to the peaks in the magnitude spectra of the four times zero-padded, cosine-apodized time domain signals. This procedure was used for analyzing both, the entire transients as well as their constituent segments. The signal-to-noise (S/N) was defined as the ratio of an observed peak apex to the *σ* of the spectral noise. For numerical experiments on the temporal stability of the ion frequency, a purely synthetic non-decaying harmonic signal with no modulations of any kind was added, at *m/z* 27130, to a 1-second-long experimental transient of a single ion of a FHV particle at *m/z* 42980 (see Supplemental Figure 3a). These two peaks, experimental and synthetic, were subsequently analyzed for stability over time.

In a first set of the computational experiments a rather short temporal window was taken ***t***_***win***_*= 64 ms*, which guaranteed baseline resolution for the interrogated peaks. The resulting smooth spectrograms (see Supplemental Figure 3a insets) were achieved via stepping with a small (relative to ***t***_***win***_) overlap of ***t***_***step***_*= 4ms*. The observed frequency modulations for both the experimental and the synthetic peaks appear to be systematic but uncorrelated. In case of the latter it is truly surprising as the artificial signal should not carry any temporal instabilities. If, however, the observed centroids are considered without the temporal information but rather as data points for histograms reporting the deviations to their expected values (see Supplemental Figure 3a insets), they appear as near-bell shaped distributions with full width at half maximum (FWHM) for the synthetic peak of ∼49ppm, and that for the experimental peak of ∼72ppm in terms of their *m/z*. Although the widths of the histograms differ in terms of ppm of their respective *m/z*’s, the peaks come from the different regions of the underlying frequency spectrum with intrinsically different granularities (i.e. for the 27130 Th peak the FT bin is 4.35 Th (∼160.44 ppm), and that for the 42980 Th is 8.68 Th (∼201.93 ppm)), and the reported spread is substantially smaller than the uncertainty imposed by the FFT grid. This all suggests that the observed mass errors are the result of distortion of spectra by noise.

Failing to observe any significant modulations in the experimental peak beyond the centroiding uncertainty (based on the behavior of the synthetic component), it is worthwhile to look into the dependency of the frequency deviation spreads on the sub-transient duration ***t***_***win***_. In the next experiment the same frequency chasing experiments were performed on the same two peaks for different ***t***_***win***_. The findings are summarized in Supplemental Figure 3b. Unlike the peak intensity uncertainty, which depends directly on the noise levels, the ***σ*** of centroiding follows the negative power trend:

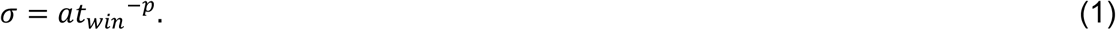

We expect both **<u>a</u>** and **<u>p</u>** to be *m/z* dependent; the investigation of this dependency is however beyond the scope of this manuscript. The observed values are *a*_42980_ = 0.025, *a*_27130_ = 0.009, and those for the exponents are *p*_42980_ = 1.45 and *p*_27130_ = 1.54. The differences scale well with the respective frequency spectrum region granularities. The overall centroiding uncertainty is well explained by noise. The apparent regularity in the observed deviations (see Supplemental Figure 3a insets) is an artifact of the finesse of sampling ***t***_***step***_. Only if the ***t***_***win***_ is extrapolated to 2 seconds (and longer) using the observed relationships (***σ***_**27130Th**_ = 0.0028 Th & ***σ***_**42980Th**_ = 0.0087 Th) the ratio between the exponential and synthetic *σ*’s become larger than one would expect based solely on the FT grid spacing suggesting that there are underlying physical phenomena (*e*.*g*. ion motion, space and image charge, *etc*.) driving the discrepancy.

The fidelity of this methodology (as that of FTMS) directly depends on the accuracy with which one can extract the frequency information (or equivalently *m/z*), where precision of peak centroiding is probably the most important step. The two biggest sources of ambiguity identified so far are the noise and the non-homogenous granularity of the Orbitrap mass spectrum. As the spacing between the data points (aka bins) in FTMS spectra tend to increase with mass, this adds to the uncertainty in accuracy of centroiding and, of course, the ambiguity brought about by noise. We focused on the effects of these two factors on the *σ* of the centroids. A set of 2D shotgun type experiments was conducted by injecting purely synthetic peaks into the experimental noisy transient across the mass range commonly used for viral particle analysis (*i*.*d*. 20-50kTh) stepping every 50 Th (to ensure thorough coverage); the S/N (defined as the ratio of a peak intensity to noise *σ*) used was ranging 6-60 with 0.2 AU step. The results are shown in Supplemental Figure 3c. As expected for sparse and baseline resolved spectra, the centroiding uncertainty increases with the noise levels, and, as was observed previously, with *m/z*. However the increase in *σ* was correlated with the mass grid spacing (a single FT bin at 20000 *m/z* = ∼0.67 Th and at 50000 *m/z* = ∼2.66 Th), *i*.*e*. 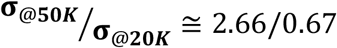.

Another parameter which affects the centroiding directly is the resolution. There are a number of factors known to affect the peak shape in FTMS (*e*.*g*. frequency instabilities, ion packet dephasing, and ion loss, *etc*.),^3–7^ yet the easiest way to manipulate the peak width is by altering the time domain signal duration. To a degree this was already done in the frequency stability analysis using different transient lengths (see Supplemental Figure 3b); here, the findings are augmented with more focused experiments. The choice was also for a 2D shotgun type experiments where, similarly, the mass was varied from 27000 Th to 27200 Th in 1 Th increments; the transient time, on the other hand, varied by power increments: 2^-5^, 2^-4^, 2^-3^, 2^-2^, and 2^-1^ of 1.024 Sec. The side-by-side comparison of the results of these *in silico* experiments with the frequency chasing experiments (for the synthetic 27130Th peak) is shown in Supplemental Figure 3d. The shotgun experiments also show the exponential dependency *σσ*_*shotgun*_ = 0.008*t*_*win*_^−1.50^. The trends are very close and made extrapolation of the “temporal evolution” experimental trend up to 0.5 sec long transients possible (see Supplemental Figure 3d). The observed discrepancy between the two is most likely caused by fewer independent data points in the chasing experiments, since it is imperative for that type of experiments to have *t*_*win*_ ≪ *t*_*transient*_.

### Experimental observations in the Orbitrap analyzer of theoretically predicted radial frequency modulations

The theoretical ion motion in the Orbitrap analyzer has been described earlier in quite some detail and combines, for a stable trajectory, rotation around as well as oscillation along the central electrode^8^. This results in a spiraling motion of the ion when injected into the orbital trap and separate terms for the frequencies of oscillations along the z (ω_z_) and the polar coordinates r (ω_r_) and φ (ω_φ_) can be formulated. The image current is detected on the split outer electrodes and the resulting signal, after a differential amplifier, is subjected to Fourier transformation. Attributed to this geometry, the final mass spectrum is calculated based on the frequencies along z. The use of oscillation frequency along z is caused by its independence on of the ion energy. This is crucial as, based on their spatial distribution in the C-trap, ions get injected with a variable amount of energy into the Orbitrap analyzer and thus occupy slightly different radii. For an ion cloud of a given *m/z*, this also means that ions will lose radial coherence rather quickly and the ions will occupy a rotating ring with a certain thickness instead of a defined point in time and space. For ensemble measurement this offers a distinct advantage since the Orbitrap analyzer can accommodate a wider range of ion energies, making it less susceptible to detrimental effects of space charging. However, for individual ions, the variable injection energy and resulting radial orbit may be problematic as the induced image current is inversely proportional to the distance of the ions to the outer electrode cups.

As mentioned in the main text, we report here the first experimental measurements of the radial frequency of ions in the Orbitrap mass analyzer. These radial harmonics are indirectly detected, as they are components of and modulate the total signal:

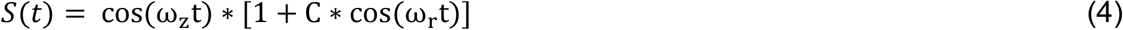

and

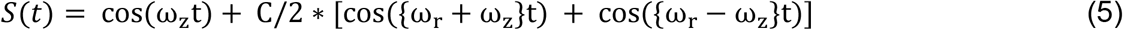

Therefore, we should find for each ion primary signal two radial frequency modulations at both higher (ω_-_=ω_r_-ω_z_) and lower (ω_+_=ω_r_+ω_z_) *m/z* (as shown in Figure 6 and Supplementary Figure 4). We observed radial frequency modulations for all theoretical expected cases like^9^: 1) highly dephased radial frequencies caused by non-circular orbits for ions with poorly matched ion energies. 2) Stable and defined radial frequencies for ions with defined orbit and well matching injection energy. 3) Varying radial frequencies for the same frequency along z. 4) Slightly reduced radial frequency for lower *m/z* species caused by an earlier arrival in the Orbitrap analyzer and thus more extended orbit.

## Supplementary Figures

**Supplemental Figure 1:**
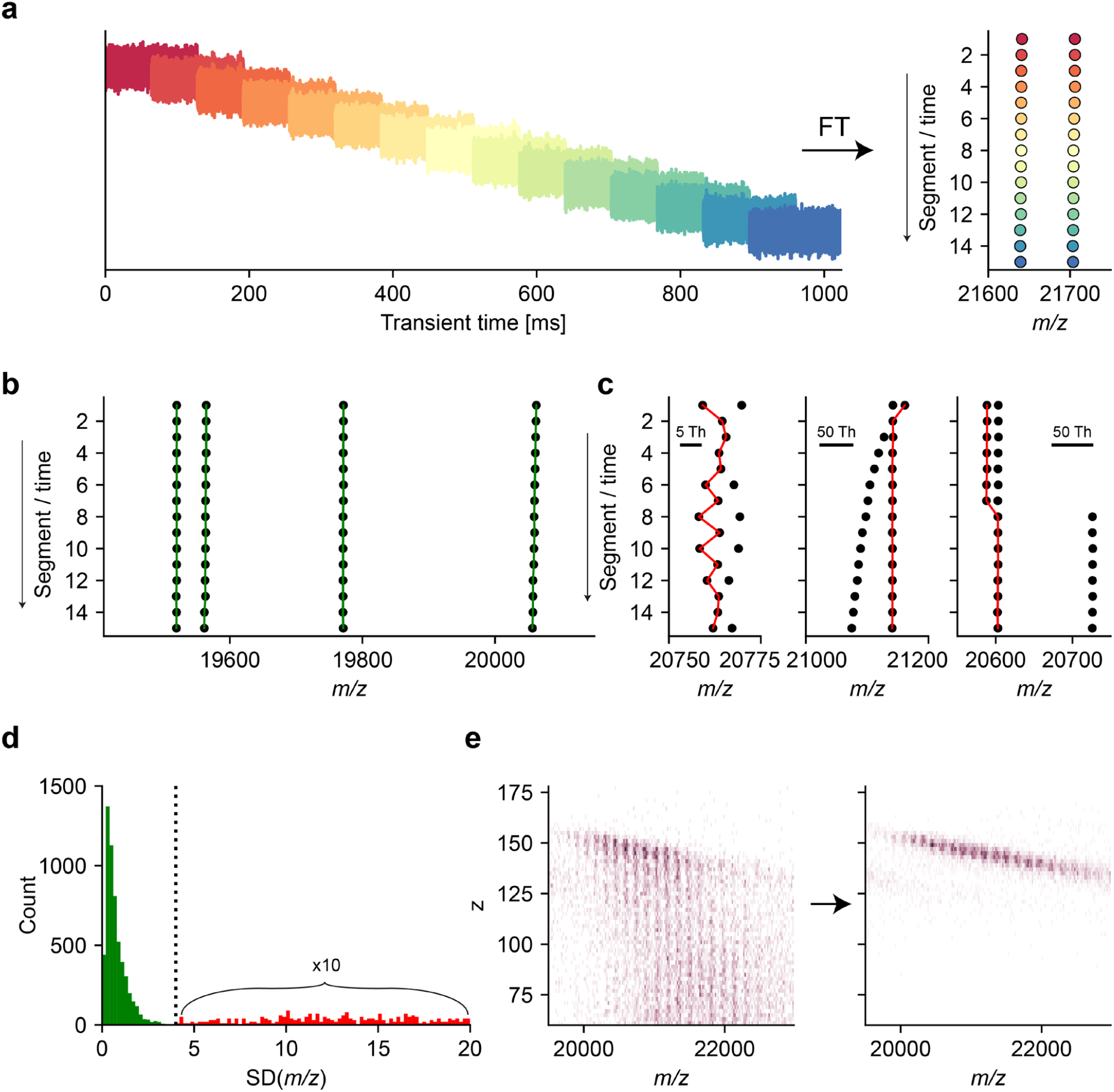
**a**, Illustration of used overlapping transient segments in the frequency chasing method and the resulting centroids per segment after FFT. **b**,**c**, Examples of ion tracing for a stable case (**b**) where all ions can be traced easily as well as a more complicated case where some ions cannot be traced due to (left to right in **c**) signals which cannot be resolved properly at the given segment lengths as well as crossing ion signals and rare charge loss events (at around 0.5% of all ion events for the given pressure setting). **d**, Histogram of the standard deviation (SD) of the average *m/z* position for traced ions where the green distribution reflects the natural pressure dependent frequency drift as well as much higher values (in red) which reflect rare occasions of falsely traced ions. The spread in SD of the successfully traced ions can also give an impression of the peak splitting prevalence as SD of truly stable ions should not exceed theoretical SD based on centroiding (>0.2Th for 128ms at 20k *m/z*), thus the majority of the peaks for the given pressure setting should exhibit some signs of peak splitting. We applied a filtering step allowing only ions under a certain threshold to be used for further analysis. **e** Illustration of the cumulative effect of frequency chasing on single particle CDMS experiments.

**Supplemental Figure 2:**
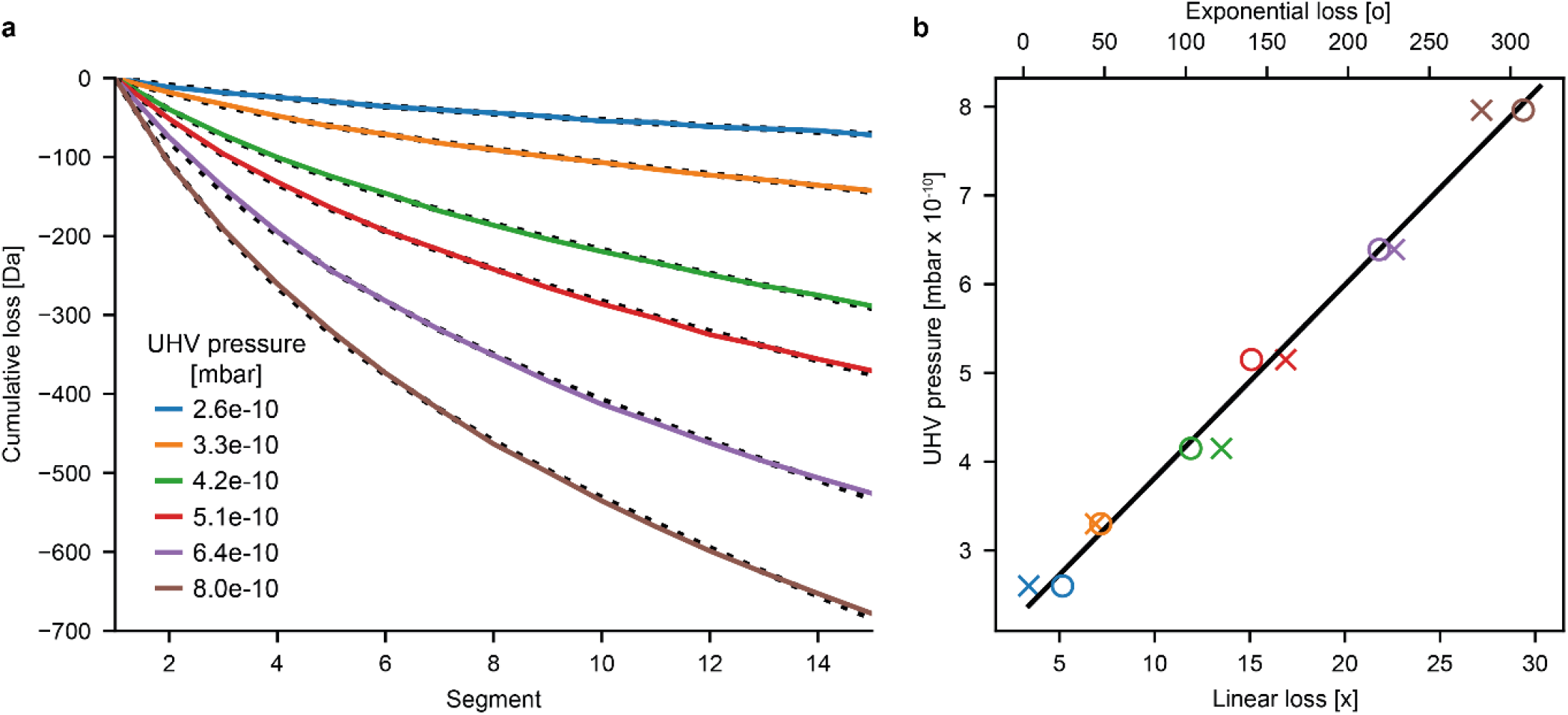
**a**, Average total neutral loss of individual ions, with respect to the first segment, plotted for a wide range of pressures. The experimental cumulative neutral loss can be described accurately with the function: *Cumulative_loss*(*Segment*) = *Exponential_loss* * e^-0.3\**Segment*^+ *Linear_loss* * *Segment*. The resulting simulations are shown as dotted lines with r^2^-values in the range of 0.998-0.9998. **b**, The corresponding values for the exponential neutral loss and linear neutral loss are plotted against the UHV pressure and show a nice linear correlation.

**Supplemental Figure 3:**
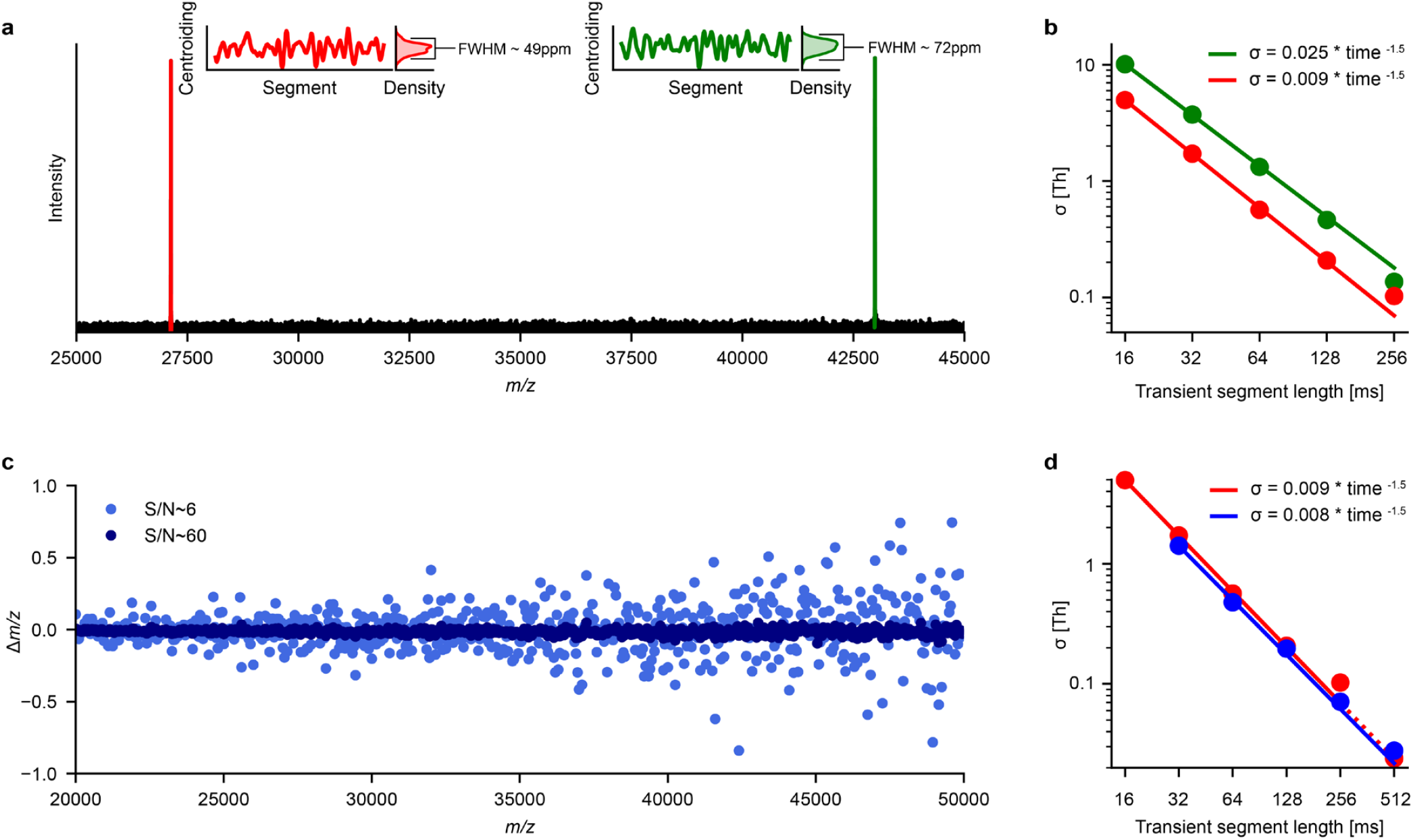
**a**, Experimental spectrum of a viral particle ion at 42980 Th with a “spiked in” synthetic species at 27130 Th. The temporal instabilities of the experimental (green) and the synthetic (red) peaks as well as the overall distribution of the observed centroids are shown as insets. **b**, The observed *m/z* stability as a function of the (sub-) transient duration. The *σ*’s of the overall distribution of the observed centroids of the experimental (green) and the synthetic (red) centroids were fitted to an exponential function. **c** The observed *m/z* uncertainty as a function of signal to noise ratio and the *m/z* values. The results of the shotgun type of computational experiments investigating the spread of the observed values (in terms of *σ*) *m/z* across the 2 kTh-5 kTh range as a function of S/N (6 to 60). **d**, The comparison of the observed *m/z* value spread in “Temporal Stability” vs “In Silico Shotgun” experiments. The observed *σ*’s of the *m/z* values of the synthetic peak at 27130 Th in the temporal evolution of the resonance frequency analysis experiments (red) and shotgun experiments (mass range is limited to 27000 Th to 27200 Th in blue) as a function of the time domain signal length. The *σ* value for the transient duration of 0.5 seconds (shown in red) is extrapolated (dotted line) using the observed relationship.

**Supplemental Figure 4:**
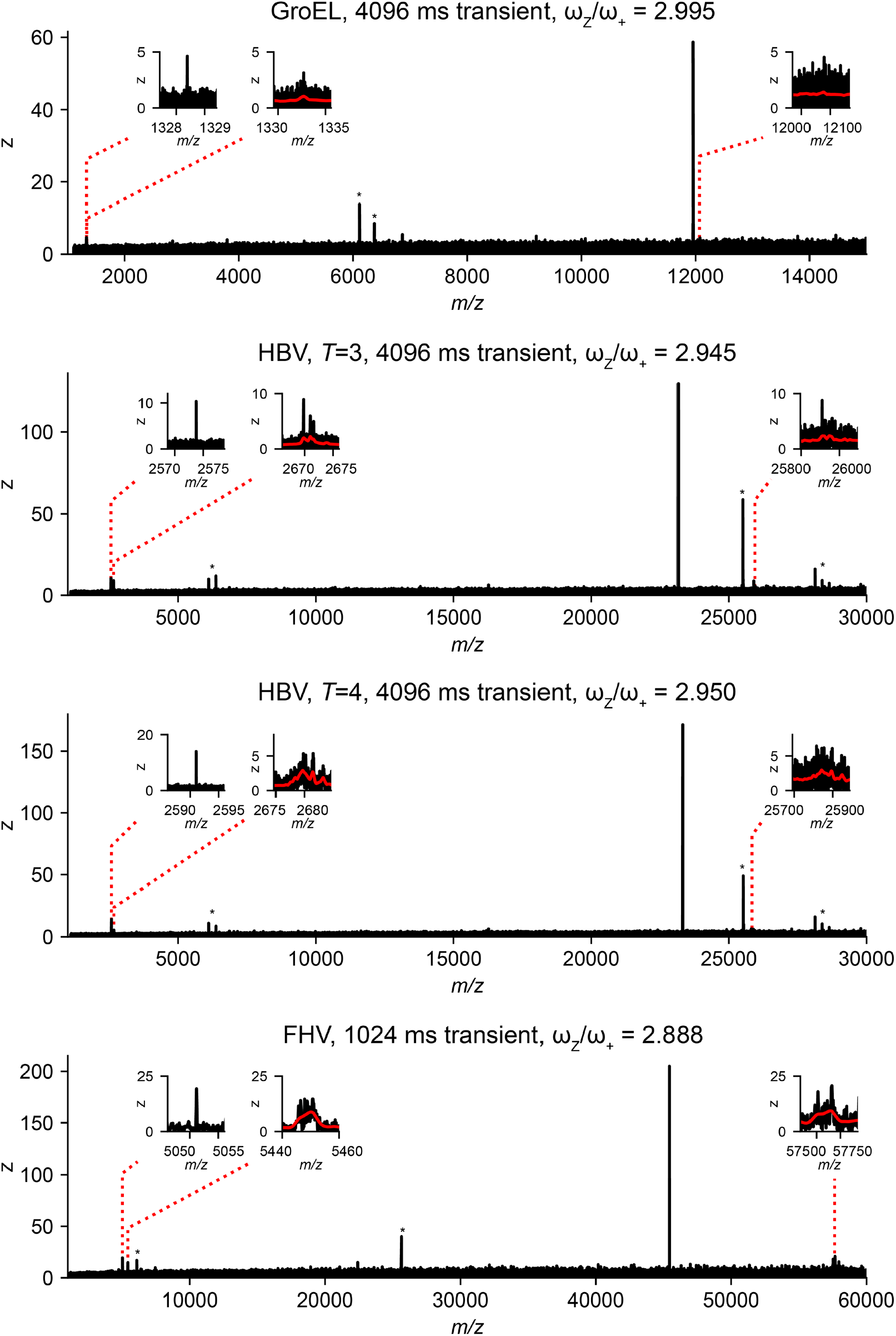
Spectra of single ion occurrences for GroEL, HBV, *T*=3 and *T*=4 as well as FHV (top to bottom). Regions where the 3^rd^ harmonic of ω_Z_, the radial frequency modulation ω_+_ and ω_-_, occur are shown as insets (left to right). The modulations for ω_-_are barely visible against the higher noise level at lower frequency and S/N for GroEL. This is contrasted by the highly charged FHV ion which modulations are clearly visible against the noise at transient duration of only 1s. We observed a continued trend to lower ω_Z_/ω_+_ for higher *m/z* values. Prominent electronic noise peaks are indicated with *.

## References

1. Leney, A. C. & Heck, A. J. R. Native Mass Spectrometry: What is in the Name? J. Am. Soc. Mass Spectrom. 28, 5–13 (2017).

2. Fenn, J. B. Electrospray wings for molecular elephants (Nobel lecture). in Angewandte Chemie - International Edition vol. 42 3871–3894 (John Wiley & Sons, Ltd, 2003).

3. Rostom, A. A. et al. Detection and selective dissociation of intact ribosomes in a mass spectrometer. Proc. Natl. Acad. Sci. 97, 5185–5190 (2002).

4. Chorev, D. S. et al. Protein assemblies ejected directly from native membranes yield complexes for mass spectrometry. Science (80-.). 362, 829–834 (2018).

5. Wörner, T. P. et al. Adeno-associated virus capsid assembly is divergent and stochastic. Nat. Commun. 12, 1642 (2021).

6. Uetrecht, C., Barbu, I. M., Shoemaker, G. K., van Duijn, E. & Heck, A. J. R. Interrogating viral capsid assembly with ion mobility–mass spectrometry. Nat. Chem. 3, 126–132 (2011).

7. van de Waterbeemd, M. et al. High-fidelity mass analysis unveils heterogeneity in intact ribosomal particles. Nat. Methods 14, 283–286 (2017).

8. Keener, J. E., Zhang, G. & Marty, M. T. Native Mass Spectrometry of Membrane Proteins. Anal. Chem. 93, 583–597 (2021).

9. Wörner, T. P., Shamorkina, T. M., Snijder, J. & Heck, A. J. R. Mass Spectrometry-Based Structural Virology. Anal. Chem. 93, 620–640 (2021).

10. Bolla, J. R., Agasid, M. T., Mehmood, S. & Robinson, C. V. Membrane Protein–Lipid Interactions Probed Using Mass Spectrometry. Annu. Rev. Biochem. 88, 85–111 (2019).

11. Zhou, M. et al. Higher-order structural characterisation of native proteins and complexes by top-down mass spectrometry. Chem. Sci. 11, 12918–12936 (2020).

12. Keifer, D. Z. & Jarrold, M. F. Single-molecule mass spectrometry. Mass Spectrom. Rev. 36, 715–733 (2017).

13. Harper, C. C., Elliott, A. G., Oltrogge, L. M., Savage, D. F. & Williams, E. R. Multiplexed Charge Detection Mass Spectrometry for High Throughput Single Ion Analysis of Large Molecules. Anal. Chem. 91, 7465 (2019).

14. Hanay, M. S. et al. Single-protein nanomechanical mass spectrometry in real time. Nat. Nanotechnol. 7, 602–608 (2012).

15. Dunbar, C. A., Callaway, H. M., Parrish, C. R. & Jarrold, M. F. Probing Antibody Binding to Canine Parvovirus with Charge Detection Mass Spectrometry. J. Am. Chem. Soc. 140, 15701–15711 (2018).

16. Brown, B. A. et al. Charge Detection Mass Spectrometry Measurements of Exosomes and other Extracellular Particles Enriched from Bovine Milk. Anal. Chem. 92, 3285–3292 (2020).

17. Dominguez-Medina, S. et al. Neutral mass spectrometry of virus capsids above 100 megadaltons with nanomechanical resonators. Science (80-.). 362, 918–922 (2018).

18. Kafader, J. O. et al. Multiplexed mass spectrometry of individual ions improves measurement of proteoforms and their complexes. Nat. Methods 17, 391–394 (2020).

19. Wörner, T. P. et al. Resolving heterogeneous macromolecular assemblies by Orbitrap-based single-particle charge detection mass spectrometry. Nat. Methods 17, 395–398 (2020).

20. Makarov, A. & Denisov, E. Dynamics of Ions of Intact Proteins in the Orbitrap Mass Analyzer. J. Am. Soc. Mass Spectrom. 20, 1486–1495 (2009).

21. Sanders, J. D. et al. Determination of Collision Cross-Sections of Protein Ions in an Orbitrap Mass Analyzer. Anal. Chem. 90, 5896–5902 (2018).

22. Kafader, J. O. et al. Measurement of Individual Ions Sharply Increases the Resolution of Orbitrap Mass Spectra of Proteins. Anal. Chem. 91, 2776–2783 (2019).

23. Rose, R. J., Damoc, E., Denisov, E., Makarov, A. & Heck, A. J. R. High-sensitivity Orbitrap mass analysis of intact macromolecular assemblies. Nat. Methods 9, 1084–6 (2012).

24. Lössl, P., Snijder, J. & Heck, A. J. R. Boundaries of Mass Resolution in Native Mass Spectrometry. J. Am. Soc. Mass Spectrom. 25, 906–917 (2014).

25. Wang, G. et al. Molecular Basis of Assembly and Activation of Complement Component C1 in Complex with Immunoglobulin G1 and Antigen. Mol. Cell 63, 135–145 (2016).

26. Diebolder, C. a et al. Complement is activated by IgG hexamers assembled at the cell surface. Science (80-.). 343, 1260– 3 (2014).

27. Terasaka, N., Azuma, Y. & Hilvert, D. Laboratory evolution of virus-like nucleocapsids from nonviral protein cages. Proc. Natl. Acad. Sci. 115, 5432–5437 (2018).

28. Sasaki, E. et al. Structure and assembly of scalable porous protein cages. Nat. Commun. 8, 14663 (2017).

29. Aizikov, K. & O’Connor, P. B. Use of the filter diagonalization method in the study of space charge related frequency modulation in Fourier transform ion cyclotron resonance mass spectrometry. J. Am. Soc. Mass Spectrom. 17, 836–843 (2006).

30. Aizikov, K., Mathur, R. & O’Connor, P. B. The spontaneous loss of coherence catastrophe in fourier transform ion cyclotron resonance mass spectrometry. J. Am. Soc. Mass Spectrom. 20, 247–256 (2009).

31. Leach, F. E. et al. Analysis of phase dependent frequency shifts in simulated FTMS transients using the filter diagonalization method. Int. J. Mass Spectrom. 325–327, 19–24 (2012).

32. Kharchenko, A., Vladimirov, G., Heeren, R. M. A. & Nikolaev, E. N. Performance of Orbitrap Mass Analyzer at Various Space Charge and Non-Ideal Field Conditions: Simulation Approach. J. Am. Soc. Mass Spectrom. 23, 977–987 (2012).

33. Abbate, A., DeCusatis, C. M. & Das, P. K. Time-Frequency Analysis of Signals. in Wavelets and Subbands 103–187 (Birkhäuser Boston, 2002). doi:10.1007/978-1-4612-0113-7_3.

34. Mann, M., Meng, C. K. & Fenn, J. B. Interpreting Mass Spectra of Multiply Charged Ions. Anal. Chem. 61, 1702–1708 (1989).

35. Uetrecht, C. et al. High-resolution mass spectrometry of viral assemblies: Molecular composition and stability of dimorphic hepatitis B virus capsids. Proc. Natl. Acad. Sci. U. S. A. 105, 9216–9220 (2008).

36. Todd, A. R., Barnes, L. F., Young, K., Zlotnick, A. & Jarrold, M. F. Higher Resolution Charge Detection Mass Spectrometry. Anal. Chem. 92, 11357– 11364 (2020).

37. Hu, Q. et al. The Orbitrap: a new mass spectrometer. J. Mass Spectrom. 40, 430– 443 (2005).

38. Makarov, A. Electrostatic Axially Harmonic Orbital Trapping: A High-Performance Technique of Mass Analysis. Anal. Chem. 72, 1156–1162 (2000).

39. Keifer, D. Z., Shinholt, D. L. & Jarrold, M. F. Charge Detection Mass Spectrometry with Almost Perfect Charge Accuracy. Anal. Chem. 87, 10330–10337 (2015).

40. Denisov, E., Damoc, E. & Makarov, A. Exploring frontiers of orbitrap performance for long transients. Int. J. Mass Spectrom. 466, 116607 (2021).

41. Fort, K. L. et al. Expanding the structural analysis capabilities on an Orbitrap-based mass spectrometer for large macromolecular complexes. Analyst 143, (2017).

42. Lange, O., Damoc, E., Wieghaus, A. & Makarov, A. Enhanced Fourier transform for Orbitrap mass spectrometry. Int. J. Mass Spectrom. 369, 16–22 (2014).

## Supplementary References

1. Posener, D. W. Precision in measuring resonance spectra. J. Magn. Reson. 14, 121– 128 (1974).

2. Chen, L., Cottrell, C. E. & Marshall, A. G. Effect of signal-to-noise ratio and number of data points upon precision in measurement of peak amplitude, position and width in fourier transform spectrometry. Chemom. Intell. Lab. Syst. 1, 51–58 (1986).

3. Makarov, A., Denisov, E. & Lange, O. Performance evaluation of a high-field orbitrap mass analyzer. J. Am. Soc. Mass Spectrom. 20, 1391–1396 (2009).

4. Gorshkov, M. V., Fornelli, L. & Tsybin, Y. O. Observation of ion coalescence in Orbitrap Fourier transform mass spectrometry. Rapid Commun. Mass Spectrom. 26, 1711–1717 (2012).

5. Kozhinov, A. N., Zhurov, K. O. & Tsybin, Y. O. Iterative Method for Mass Spectra Recalibration via Empirical Estimation of the Mass Calibration Function for Fourier Transform Mass Spectrometry-Based Petroleomics. Anal. Chem. 85, 6437–6445 (2013).

6. Ledford, E. B., Rempel, D. L. & Gross, M. L. Space charge effects in Fourier transform mass spectrometry. Mass calibration. Anal. Chem. 56, 2744–8 (1984).

7. Jeffries, J. B., Barlow, S. E. & Dunn, G. H. Theory of space-charge shift of ion cyclotron resonance frequencies. Int. J. Mass Spectrom. Ion Process. 54, 169–187 (1983).

8. Makarov, A. Electrostatic Axially Harmonic Orbital Trapping: A High-Performance Technique of Mass Analysis. Anal. Chem. 72, 1156–1162 (2000).

9. Hu, Q. et al. The Orbitrap: a new mass spectrometer. J. Mass Spectrom. 40, 430–443 (2005).

